# Ten numbers from the Laplace–Beltrami spectrum facilitate training-free classification of protein structures based on surface shape

**DOI:** 10.64898/2026.09.10.750378

**Authors:** M. Fernández-Giro, J. Emonts, B. Berkels, J. F. Buyel

## Abstract

The comparison of protein structures is necessary to determine molecular functions and evolutionary relationships and can also facilitate drug design and the prediction of separation options. The number of available protein structures is rapidly increasing, driven by experimental determination but also prediction methods such as AlphaFold. In this context, traditional structure comparison based on atomic superposition becomes computationally inefficient. To enable scalable analysis and surface shape comparison, alternative approaches represent protein structures using compact, fixed-length vectors known as descriptors. Current vectors often contain hundreds of entries, such as three-dimensional Zernike descriptors composed of 121 entries, or combinations of molecular and geometric properties. Here, we propose to use the Laplace–Beltrami spectrum, a mathematical representation of object surfaces derived from the eigenvalues of the Laplace–Beltrami operator, as an efficient option to capture protein structural properties and facilitate structural classification. Specifically, protein surfaces are encoded using only the first 10 non-zero eigenvalues of this spectrum. We used the SHREC 2025 dataset as a benchmark and achieved 85.8% accuracy on the original 97 protein structure classes and 97.7% accuracy on the homology-grouped 45 classes. Our method is fast, requiring ∼75 min computation time for the 11,555 protein surface meshes of the SHREC 2025 dataset on a conventional AMD Ryzen 7 5700X 8-Core processor with 32 GB RAM. The approach can easily be applied to large protein structure databases because it is readily parallelizable and allows the incorporation of other descriptors, including surface features such as charge. This rapid screening tool can therefore be used to identify proteins with related surface shapes independent of sequence homology.

## 1. Introduction

Proteins fulfil a vast range of biological functions enabled by their diverse surface properties and three-dimensional (3D) architectures known as folds. Although the number of folds is limited, indicating conservation across evolutionary processes, there is immense combinatorial diversity in amino acid sequences (Holm and Sander 1996; Levitt 2009; Ingles-Prieto et al. 2013). This can hinder the identification and classification of protein structures based on the primary amino acid sequence. Traditional approaches for protein structural classification, such as Structural Classification of Proteins (SCOP) (Fox et al. 2015; Andreeva et al. 2020), Class, Architecture, Topology, and Homologous superfamily (CATH) (Sillitoe et al. 2021), and the Evolutionary Classification of Protein Domains (ECOD) (Cheng et al. 2014), rely on the hierarchical organization of proteins based on sequence similarity, secondary structures, and overall fold topology. These frameworks have defined the important principles of structure–function relationships, showing that proteins with low sequence identity can nevertheless adopt highly similar 3D architectures. At their core, such classifications rely on atomic-level structural superposition and manual or semi-automated curation, making them computationally intensive and difficult to scale to the rapidly expanding universe of experimentally resolved and computationally predicted protein structures. Accordingly, there is an increasing need for fast, geometry-driven methods enabling the reliable comparison and classification of protein structures even with little sequence homology. In this context, the SHape REtrieval Contest (SHREC) series has emerged as a valuable benchmarking platform for protein structure classification (Mavridis et al. 2010, Song et al. 2017, Yacoub et al. 2025). Sequence-independent structure classification tools are useful because convergent evolution can result in similar protein structures even in the absence of a common ancestor. For example, enzyme active sites and ligand-binding domains are often defined by key conserved residues embedded within distinct geometries (Galperin and Koonin 2012; Shafee et al. 2017). Accordingly, shape and surface comparison methods have the potential to uncover functional analogies that are difficult to detect by sequence or fold homology. In this context, 3D Zernike descriptors (Sael et al. 2008; La et al. 2009) represent a protein structure compactly through a series expansion in orthogonal 3D polynomials, typically normalized to achieve rotation invariance and scaled to the unit sphere. Although effective for capturing global volumetric shape, these descriptors are extrinsic in nature and thus sensitive to isometric deformations such as bending.

Here, we describe and compare protein shapes using the Laplace–Beltrami spectrum (LBS), calculated by subjecting a discretized protein surface mesh to the Laplace–Beltrami operator (LBO). The LBO is a generalization of the Laplace operator, representing the divergent of the gradient of a scalar function in Euclidian space, to manifolds (Buser 2010). A manifold can be thought of as a potentially curved object that appears locally flat, like the surface of the earth perceived by a human, a concept that applies to any sphere in general. As such, the LBO is a central construct in spectral geometry, capturing the intrinsic geometry of a manifold independently of its embedding in 3D space. In particular, two-dimensional manifolds represent surfaces such as those of proteins. The eigenvalues of the LBO (λ_k_), i.e., the LBS, are often described as the “shape-DNA” of an object (Reuter et al. 2006), and encode geometric information across multiple scales, from global structure to local curvature, thus providing an isometry-invariant shape signature. This approach has been successful in computer vision for object recognition and, more recently, for small molecules in virtual screening contexts (Seddon et al. 2019), but its application to macromolecular protein surfaces has yet to be systematically explored. Here, we hypothesize that proteins with divergent sequences, but conserved surface morphologies, will exhibit a similar LBS, enabling the detection of remote structural analogs and signatures of functional convergence. Leveraging the large dataset provided by the SHREC 2025 challenge (Yacoub et al. 2025), we assess the ability of these spectral surface descriptors to discriminate and classify protein structures based solely on surface geometry.

## 2. Materials and methods

### 2.1 Protein surface mesh generation

Protein structure files (Figure 1A; Table S1) were converted from *.pdb to *.xyzr format using values for atomic radii given by AMBER99 (Wang et al. 2000), CHARMM22 (Mackerell et al. 2004) or Swanson (Swanson et al. 2007) force fields (Figure 1B), taken from *.dat files in the Adaptive Poisson-Boltzmann Solver (APBS) software (Jurrus et al. 2018) (https://github.com/Electrostatics/pdb2pqr). Probes of 0.11–0.19 nm (1.1–1.9 Å; ∼0.75–1.25 times the size of a water molecule) were used to create and triangulate solvent-excluded surfaces (Connolly 1983) using the Michael Sanner Molecular Surface (MSMS) script (Sanner et al. 1996). The selected triangular mesh densities were of 100–500 vertices nm^-2^ (1–5 vertices Å^-2^). Additionally, a density of 150 vertices nm^-2^ (1.5 vertices Å^-2^) was selected for compatibility with the SHREC 2025 benchmark dataset (Yacoub et al. 2025) (Figure 1C). For proteins in the SHREC 2025 dataset, surface meshes were based on the solvent-excluded molecular surfaces calculated in NanoShaper (Decherchi et al. 2019). Disconnected, internal mesh artifacts were eliminated before LBS computation by removing all but the largest mesh from each surface file.

**Figure 1.**
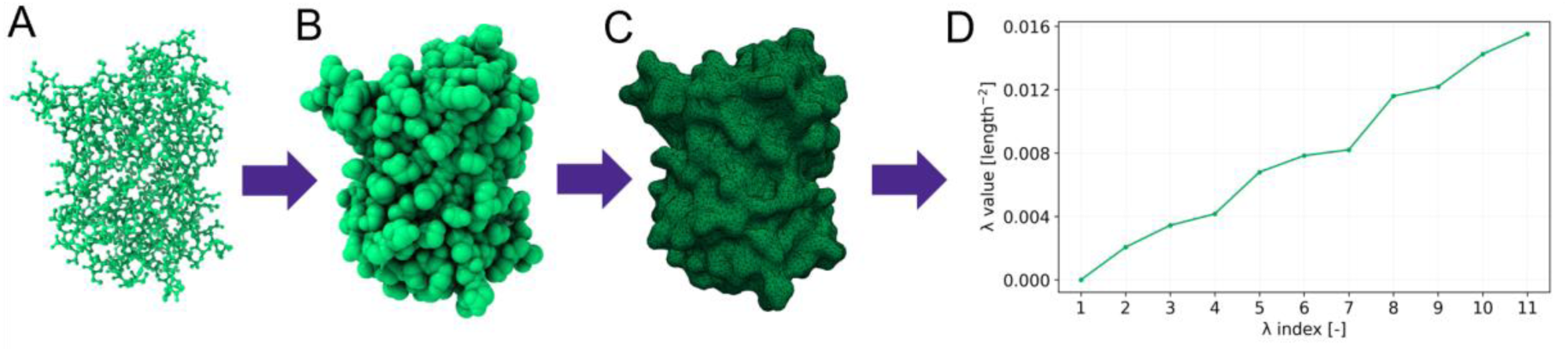
Schematic workflow for the calculation of a protein Laplace–Beltrami spectrum. A. Atomic coordinates taken from PDB files. Here, green fluorescent protein (GFP; PDB ID 1ema) was used as an example. B. Radii given by the AMBER99 force field (Wang et al. 2000) were assigned to atomic coordinates in *.xyzr files. C. Using the Michael Sanner Molecular Surface (MSMS) script (Sanner et al. 1996), surfaces were converted to triangular meshes using a probe radius of 0.15 nm and a resolution of 150 vertices nm^-2^ (1.5 vertices Å^-2^). D. Laplace–Beltrami spectrum of GFP (1ema) from λ_1_ to λ_11_. Note that λ_1_ is always zero (in our setting of connected meshes) and thus does not hold geometric information. See Figure S1 for more details.

### 2.2 Laplace–Beltrami spectrum computation

The LBS was calculated by solving the eigenvalue problem (Equation 1).

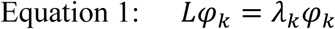

where *L* is the LBO, and φ_k_ and λ_k_ are the *k*^th^ eigenfunction and eigenvalue of the LBO respectively, which can be rewritten as Equation 2 and re-stated as Equation 3.

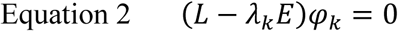

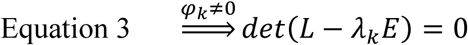

where *E* is the identity.

A piecewise linear finite element method (FEM) was used to obtain a discretization of the LBO to be applied to the triangular protein surface meshes and to compute the first N eigenvalues, where N was in the 11–100 range. The FEM was used as implemented in Python (Seddon et al. 2019) on a single AMD Ryzen 7 5700X 8-Core Processor, with 32 GB of RAM, running Windows 10 and Python 3.13. LBO eigenvalues were either used as-is or as eigenvalues of an LBO scaled by multiplying each λ_k_ by the corresponding protein surface mesh area (i.e., normalized for protein surface area).

### 2.3 Using the SHREC 2025 dataset for benchmarking

We used the SHREC 2025 dataset (Yacoub et al. 2025) to asses protein shape classification. Its solvent-excluded surface meshes for 11,555 proteins were derived from experimentally determined structures. A ground truth of 97 imbalanced classes was established by grouping proteins with >98% sequence identity. A second reference classification was built by assigning meshes of proteins with >90% sequence identity into 45 classes (Figure S2). The original dataset was split into 80% training and 20% test surface meshes on a per-class basis. In addition to this static training–test assignment, we re-shuffled the training and test set assignments of surface meshes 10 times, always maintaining an 80-to-20% distribution. Information about the biological function of individual proteins was retrieved from the Protein Data Bank (PDB) based on the protein identifiers provided in the SHREC 2025 dataset.

### 2.4 Surface classification based on the Laplace–Beltrami spectrum

In our LBS approach, each surface in the test set was assigned to a structure class based on its 1-Nearest-Neighbor in the training set (Figure 2). The distance (i.e., shape difference) between surfaces corresponds to the Manhattan distance (Equation 4), defined as the sum of absolute differences between the elements of two vectors of equal length. Here, the selected vectors consisted of the first 11 eigenvalues (λ_1_–λ_11_; note that λ_1_ is always 0 and could thus be excluded but we decided to keep it in the calculation for simplicity of formulation) of the LBS of each surface mesh. Specifically, the metric was calculated for each pair of test surfaces *n* and training surfaces *m*, giving rise to an *n* × *m* matrix.

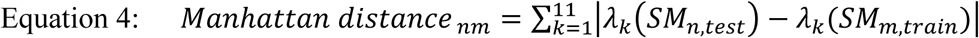

**Figure 2.**
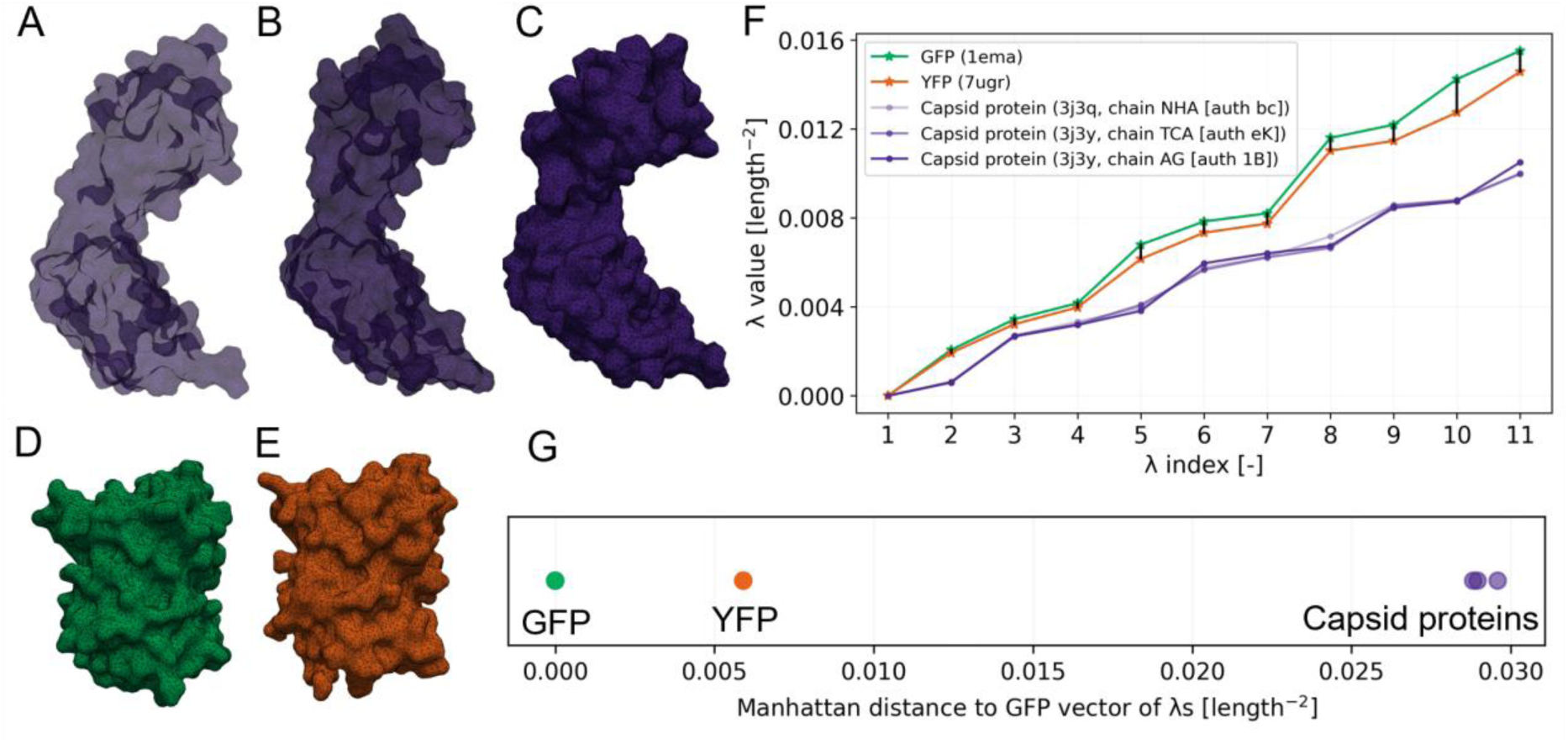
Protein surface classification workflow using the Manhattan distance metric. A–E. Surface meshes of two different protein classes belonging to either capsid protein p24 from human immunodeficiency virus 1 (HIV-1) (A, PDB ID 3j3q, chain NHA [auth bc]; B, PDB ID 3j3y, chain TCA [auth eK]; C, PDB ID 3j3y, chain AG [auth 1B]) or fluorescent proteins GFP (D, PDB ID 1ema) and YFP (E, PDB ID 7ugr). F. LBO eigenvalues (λ_1_–λ_11_) calculated for meshes shown in A–E. As an example, differences in eigenvalues between GFP (D) and YFP (E) are shown as vertical black lines. The sum of these lines is the Manhattan distance (Equation 4). G. Plot of Manhattan distances between example surface meshes (A–E) using GFP (D) as the reference point.

where *k* is the eigenvalue index, SM_n,test_ is the surface mesh of the *n*^th^ test set surface to be classified, and SM_m,train_ is the *m*^th^ surface mesh of the training set. In addition, the Euclidean distance was computed and used for comparison (Equation 5). It is defined as the square root of the sum of squared differences between corresponding elements.

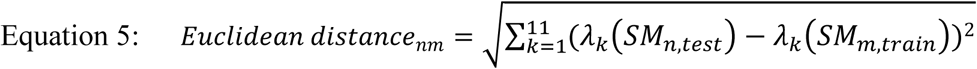

The classification performance was assessed based on the accuracy, balanced accuracy, F1 score, precision and recall (Yacoub et al. 2025) (Equations 6–10).

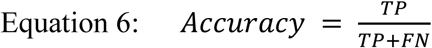

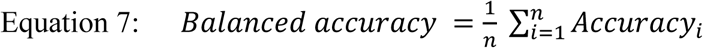

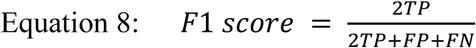

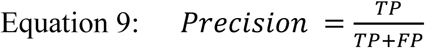

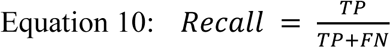

where *TP* is the number of true positives (assigned class *i*, true class *i*), *FP* the number of false positives (assigned class *i*, true class not *i*), *FN* the number of false negatives (not assigned class *i*, true class *i*) and *n* the number of classes. Note that in this setting of a single-label multiclass supervised classification approach, the recall is equal to the accuracy.

### 2.5 Alignment-based protein structure similarity

The Combinatorial Extension (CE) algorithm (Shindyalov and Bourne 1998) implemented in the Pairwise Structure Alignment tool of the PDB (https://www.rcsb.org/alignment) (Bittrich et al. 2024) with default parameter settings was used as the reference method to evaluate structural protein similarity. This was based on the root mean square deviation (RMSD) and the TM-score (Xu and Zhang 2010) using green fluorescent protein (GFP) and yellow fluorescent protein (YFP) from *Aequorea victoria* (1ema, 7ugr; 90% sequence identity) as well as GFP from *Pontellina plumata* (2g3o, chain A; with 19% sequence identity compared to 1ema) for comparison. For sequence-dependent alignments the Smith-Waterman 3D algorithm (Smith and Waterman 1981) was selected as the alignment method in https://www.rcsb.org/alignment.

## 3. Results and discussion

### 3.1 Impact of protein surface mesh preparation on the Laplace–Beltrami spectrum

The LBO eigenvalues (λ_k_) describe protein surface curvature and have the dimension of an inverse area (e.g., nm^-2^). Accordingly, they scale inversely with the surface area of the described object, in this case a protein, and we reasoned that protein surface mesh generation settings can affect and potentially distort their values. Therefore, we quantified the impact of atomic radii, mesh densities and solvent probe radii on three representative levels in MSMS (Sanner et al. 1996) when generating protein surface meshes from atomic coordinates (Tables S2 and S3). We found that the average relative standard deviation (RSD) of the first 11 eigenvalues was 1.12% and 2.51% when atomic radii and mesh density were varied, respectively (Figure S3; Tables S2 and S3). Also, the RSD was only 8.07% when we tested a variation in probe radii of about ±25%. The latter represented a substantial change given the assertiveness of the average water molecule radius (Graziano 2004). We attributed the small differences between spectra primarily to discretization effects arising from approximating the continuous molecular surface using a triangulated mesh. Accordingly, when normalizing λ_k_ for the surface mesh area, the average RSD for these eigenvalues decreased to 1.06%, 0.36% and 2.09%, respectively, as expected based on the inverse area-dependent scaling of the LBS. Even so, we continued to use the non-normalized eigenvalues for all subsequent tasks because this conserved information about protein size and we considered the associated RSD as acceptable given that the average relative difference between two closely related structures such as GFP (PDB ID 1ema) and YFP (PDB ID 7ugr) was 1.4% when using the same settings during mesh generation (Figure S3D). Furthermore, we decided to use AMBER99 atomic radii, a solvent probe radius of 0.15 nm, and a mesh density of 150 vertices nm^-2^ (1.5 vertices Å^-2^) to ensure compatibility with our other modeling work, to accommodate the most common solvent (i.e., water), and to ensure compatibility with meshes provided in the SHREC 2025 dataset (Yacoub et al. 2025), respectively. Likewise, we continued with a mesh density of 150 vertices nm^-2^, as used in the latter dataset, because it had a small impact on performance given that λ_k_ values are calculated only once for each protein and computation time increased in a near-linear manner with increasing mesh density (R^2^ = 0.98). Using these settings, the calculation of the first 11 and 100 Laplace–Beltrami eigenpairs (eigenvalues and associated eigenfunctions) took 75 min and 330 min, respectively, for all 11,555 proteins in the SHREC 2025 dataset using a conventional desktop computer.

### 3.2 Surface mesh quality control and artifact handling

We noticed that the SHREC 2025 dataset (Yacoub et al. 2025) contained entries where small, apparently artificial, close-to-spherical meshes appeared inside the expected protein surface mesh, as well as larger cavities with irregular shapes (Figure S4). Specifically, 6641 of the 11,555 dataset files contained more than one surface mesh. These artifacts undermined the numerical stability of the LBS computation and generated multiple zero-value eigenvalues. Because these artifact meshes only appeared in files of the SHREC 2025 dataset and not in surfaces processed using MSMS, we suspect that most artifacts arose during mesh generation in Nanoshaper (Decherchi et al. 2019), which can detect cavities in proteins and can include the former as disconnected mesh components. The disconnected meshes were therefore filtered out before the LBS calculation, which requires a single continuous manifold (Seddon et al. 2019).

Of greater concern was that at least 11 of the 6641 multi-mesh files contained discontinuous or disjunct surface meshes representing several of the protein’s residues, three of which were removed in an updated version of the dataset (https://gitlab.com/ycbtaher/shrec2025; update on 2025-10-28; Figure S5A–C), because this indicated substantially compromised structural information. Such files were readily identifiable within their protein structure class due to deviations in their LBS (see Section 3.6) or by the relative mesh area difference between the largest and second largest meshes. We retained only the largest partial surface mesh in these files for structure classification. Accordingly, we elected to exclude the remaining (at least) eight disjunct meshes from the current dataset and recommend the same approach for all future reference datasets to ensure reliable protein surface classification results. Likewise, single-mesh files coming from structures with missing residues were distinguishable within their classes by the smaller surface area, which in turn affects the LBS (Figure S5).

In contrast to the description of the SHREC 2025 dataset, we found that the latter contained 13 meshes with only 17–28 amino acid residues (PDB IDs 6hoh, 7lt6, 7brt and 7lsw in classes 73 and 85), lacking corresponding meshes of the full-length proteins (Table S4). We nevertheless used these meshes in the subsequent classification to ensure comparability with previous results. However, in our opinion, such files should be excluded from reference datasets for classification tasks because the absence of proper reference structures can result in modeling artifacts and potentially overfitting.

### 3.3 Selection of Laplace–Beltrami operator eigenvalues

The maximum possible number (*k*) of eigenvalues (λ) of the LBO corresponds to the number of vertices in a protein surface mesh. Using the same set of λ_k_ for all proteins facilitates structural comparison because the resulting shape descriptor vectors have the same size (i.e., an identical number of elements). Whereas comparison is faster for a smaller number of λ_k_ per surface mesh, classification accuracy can decrease because some surface features may not be well represented in the LBS. Accordingly, a suitable range of eigenvalues must be identified for the LBS.

To do so, we plotted the classification accuracy against the LBS length of continuous sets of λ values in the range λ_1_–λ_1_ to λ_1_–λ_100_ (Figure 3). We limited the range to the first 100 λ_k_, thus ensuring acceptable computation times of ∼1.7 s per surface mesh and because others have reported that fewer than 20 eigenvalues are often sufficient to describe the global shape of an object (Reuter et al. 2006). Here, we found that λ_1_–λ_11_ achieved the highest classification accuracy of 85.9 ± 0.3% (n = 11 splits; ± standard deviation) for the 97 classes defined in the SHREC 2025 dataset (see Section 3.4).

**Figure 3.**
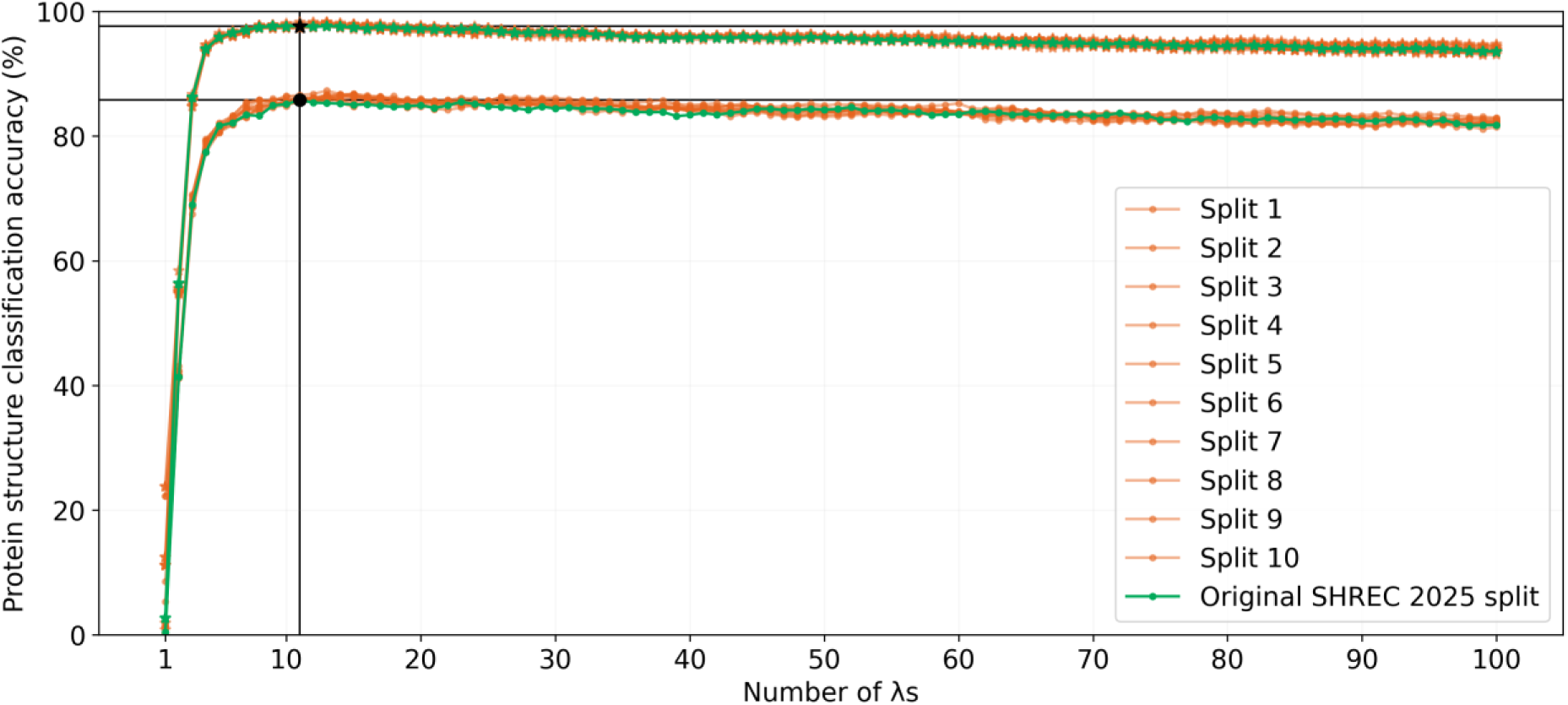
Protein structure classification accuracy achieved with the Laplace–Beltrami spectrum (LBS) up to λ_100_ for the SHREC 2025 dataset based on a 1-Nearest Neighbor algorithm. Dots indicate classification accuracy using the original 97 classes defined in the dataset. Stars indicate classification accuracy using a reduced set of 45 homologous classes as defined in the SHREC 2025 dataset. Accuracy (Equation 6) was calculated for the classification of test set surface meshes using the indicated number for λ_k_. For example, the black dot indicates the accuracy of 85.8% achieved using the first 11 eigenvalues of the LBO (λ_1_ to λ_11_). The green data series indicates results for the original data split into training and test sets, whereas the orange data series indicates results for 10 alternative splits of the data.

Interestingly, increasing the number of eigenvalues beyond 11 resulted in a reduction of the classification accuracy. This may indicate that minute and probably irrelevant local surface attributes (e.g., λ_k_ where k > 10) increasingly outweighed the (in the current classification context) more important global shape features. Specifically, each eigenvalue has a characteristic length scale proportional to k^−0.5^, with the first eigenvalues capturing the coarse protein shape and topological configurations whereas eigenvalues late in the spectrum describe local shape features (Reuter et al. 2006).

We acknowledge that it is possible to apply feature selection methods on the LBS to either reduce the number of eigenvalues in the shape descriptor vector even further or identify sets of individual eigenvalues that improve the classification accuracy. For example, using only λ_1_–λ_5_ when seeking remote structural analogs or focusing on λ_k_ >> 10 to distinguish closely related proteins can also be helpful. In this context, selecting discontinuous sets of λ_k_ may introduce additional dependencies on the training data and hence potential biases, which is why we used the continuous set of λ_1_–λ_11_ for all further tasks.

### 3.4 Classification performance on the SHREC 2025 dataset

The SHREC 2025 dataset had 9244 labeled surfaces for training and 2311 unlabeled surfaces to be assigned to the 97 original or 45 merged classes. Although electrostatic properties were available in the data (Yacoub et al. 2025), we deliberately restricted our analysis to the surface meshes, allowing us specifically to evaluate the shape classification power of the LBS, excluding its eigenfunctions.

For the classification process, protein features, here the LBS, need to be compared so similarities in shape can be identified. Many comparison metrics place an emphasis on avoiding few but extreme deviations between two samples like protein structures by first squaring differences between the sample features. This renders these approaches sensitive to extreme values (also known as outliers) (Willmott and Matsuura 2006) as is the case for RMSD used in protein structure alignment or the Euclidean distance to compare feature vectors. Because the absolute values and standard deviation of λ_k_ increase monotonically with *k* (e.g., in the present dataset, λ_11_ has 7.6 times the standard deviation of λ_2_), using one of the above approaches would have placed artificial weight on λ values with large *k* (i.e., local protein surface features). Therefore, we used the Manhattan distance metric (Section 2.4) to avoid this bias, which indeed improved the performance compared to the Euclidean distance (Table 1).

**Table 1.**
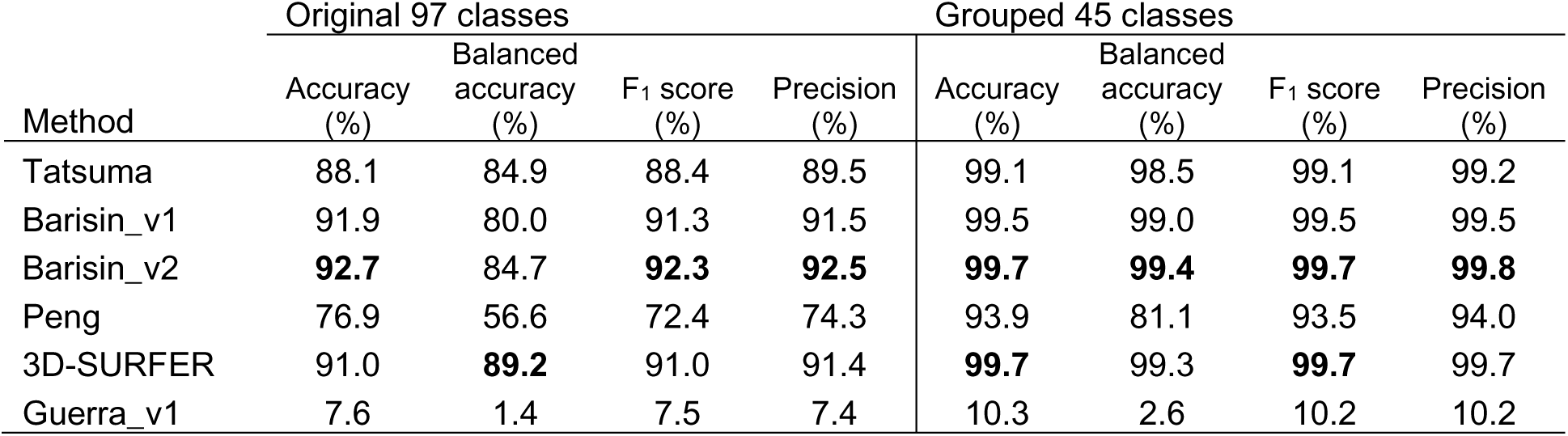
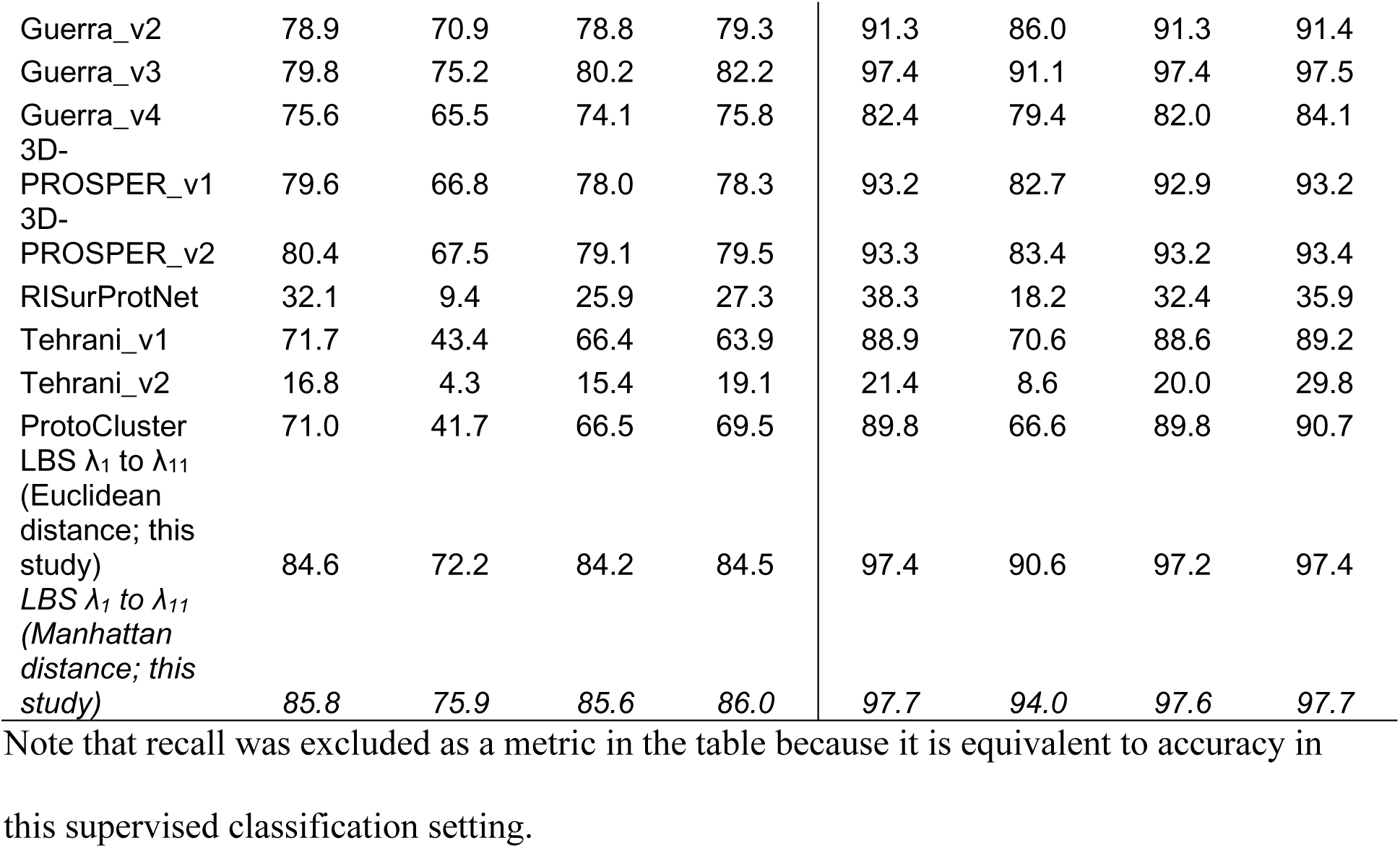
Performance metrics for protein structure classification approaches. Best values per metric are shown in bold, and the current best LBS approach is italicized.

Using λ_1_–λ_11_, the classification accuracy (here equal to the recall metric) on the initial 97 classes was 85.8%, with 1983 protein surface meshes correctly classified (assigned to the class in which they belong) and 328 misclassified (assigned to a different class than where they belong, also called false negatives as defined in Section 2.4). Specifically, 88 of the 97 classes contained test set proteins, and of these, 38 classes had a per-class (i.e., balanced) accuracy of >90% (Figure 4). Precision and F_1_ score (i.e., the harmonic mean of the precision and recall) were consistently above 0.85 (Table 1).

**Figure 4.**
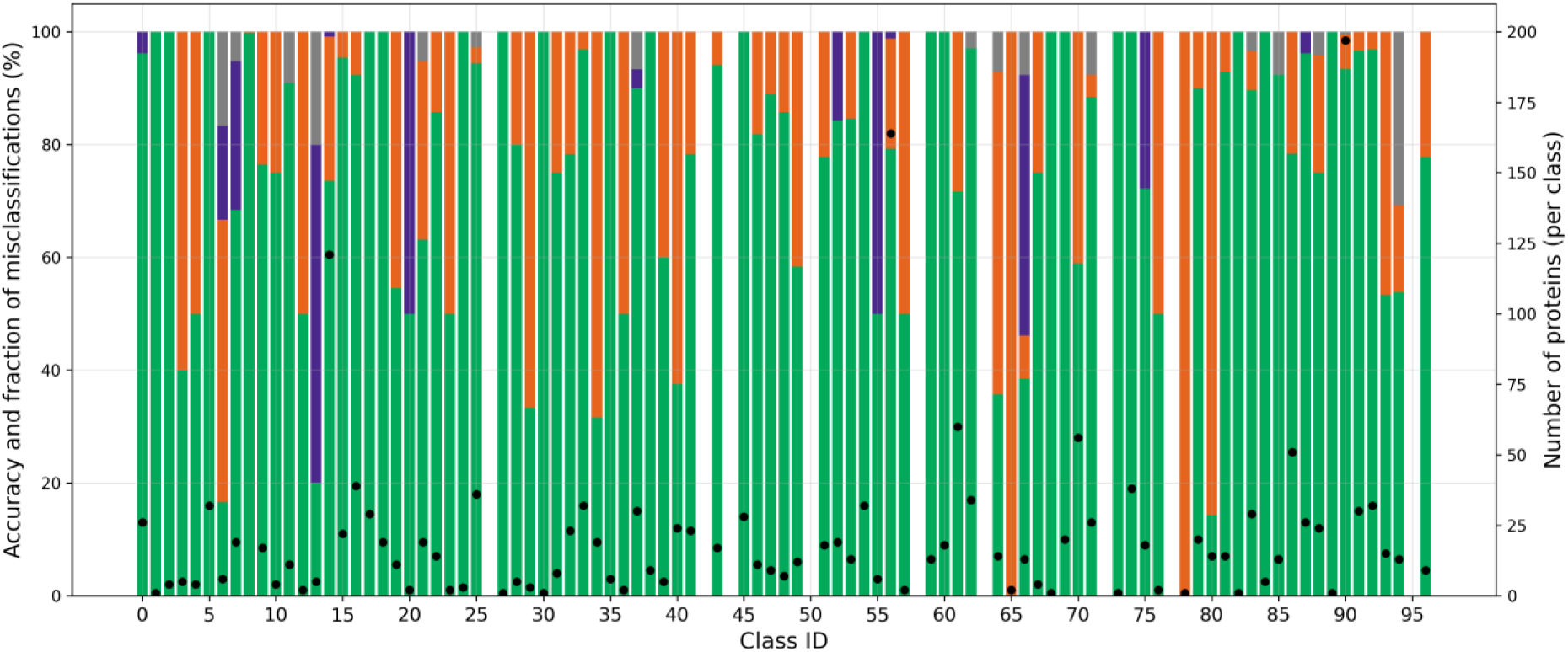
Classification performance for each of the 97 protein classes as defined in the SHREC 2025 dataset and ordered by their identifier on the *x*-axis. Green column parts indicate the fraction of correctly assigned proteins, orange marks the fraction of surfaces misclassified as homologous classes, purple represents the fraction of non-homologous but functionally related surfaces, and gray is the fraction of functionally unrelated misclassifications in non-homologous classes. Black dots indicate the number of test set proteins per class, with class 8 containing 514 proteins (out of the axis range). Nine classes did not contain any test set protein, shown as empty white columns.

### 3.5 Analysis of suspected misclassifications I – functionally-related proteins

Of the 14.2% of all test meshes that were apparently misclassified, most (11.9% of all test meshes) were assigned to a homologous class among the 97 classes. Accordingly, after aggregating the original 97 into 45 homologous classes as described above (Yacoub et al. 2025), the classification accuracy increased to 97.7% with only 54 misclassifications found in non-homologous classes (Figure 4, Table S5).

Interestingly, 33 (∼61%) of these remaining misclassifications were functionally related to the class to which they had been misassigned. Specifically, 22 Nearest Neighbor pairs of surface meshes corresponded to chaperonin subunits, three correspond to histones and one was found between proteasome subunits. For example, some pairs of these proteins shared overall sequence identities of 21%, 29% and 22%, respectively (Figure 5). Accordingly, they can be regarded as distant sequence homologs but, due to their function, close structural homologs (Willison 2018; Park et al. 2024), which was easily identified by the LBS. The same applied to seven ribosomal proteins (Table S5).

**Figure 5.**
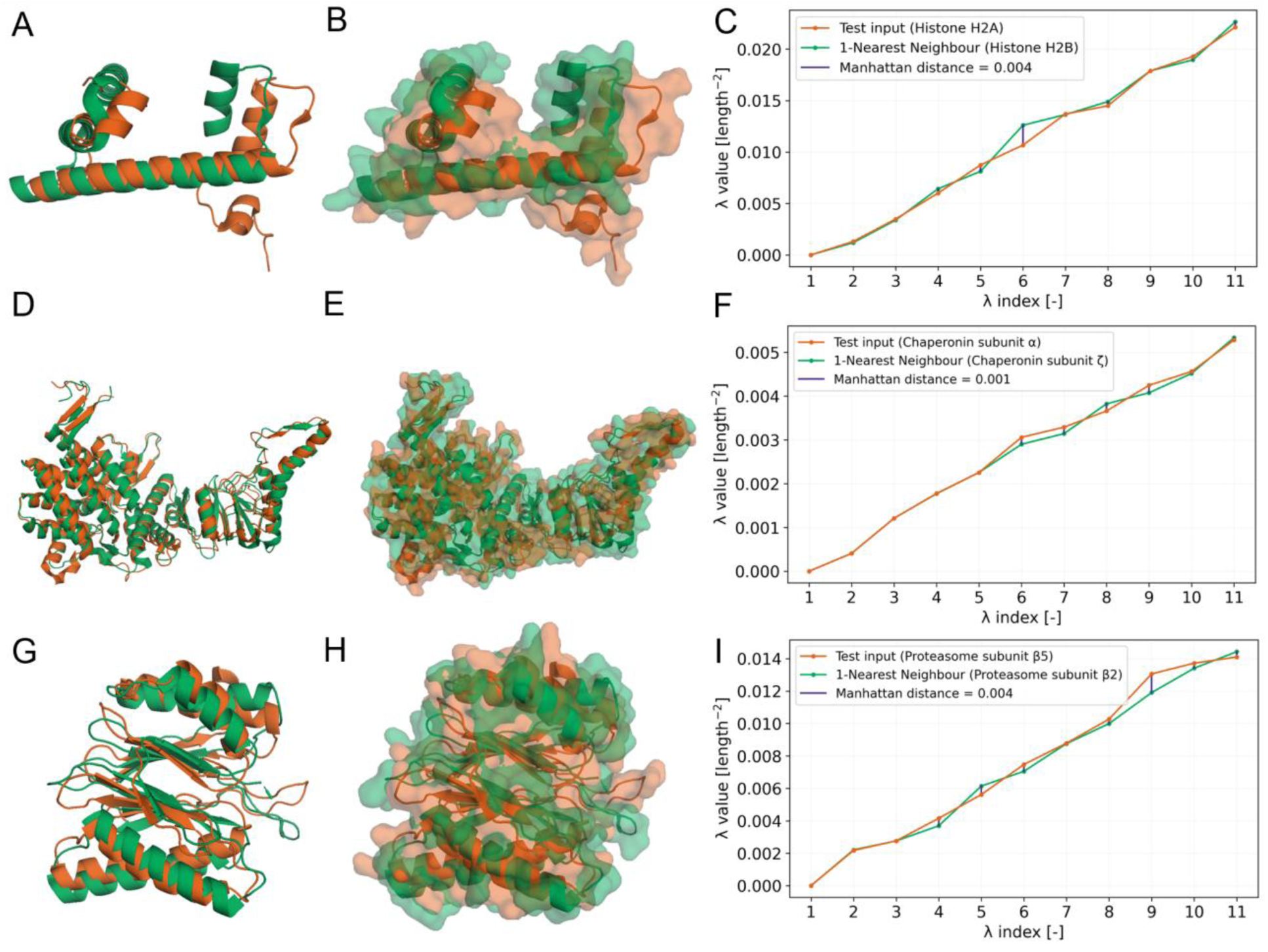
Analysis of surface mesh misclassification to non-homologous but functionally related protein classes. Ribbon representation (A), surface (B) and Laplace–Beltrami spectrum (LBS) (C) of a histone 2A (orange, test ID 637, test revealed ID 4kha_2:B:B, class 14) and its Nearest Neighbor, a histone 2B (green, train ID 3kxb_4_D_D_model1, class 93), discerned by the smallest Manhattan distance (λ_1_–λ_11_). This pair shares 21% sequence identity and 2.8 Å root mean square deviation (RMSD) over 70 aligned residues. Both proteins originate from *Xenopus laevis*. Ribbon representation (D), surface (E) and LBS (F) of a subunit α of the chaperonin complex (orange, test ID 1321, test revealed ID 8i1u_1:I:I_model1, class 7) and its Nearest Neighbor, a subunit ζ of the chaperonin complex (green, train ID 8i1u_6_N_N_model1, class 75), discerned by the smallest Manhattan distance (λ_1_–λ_11_). This pair shares 29% sequence identity and 1.9 Å RMSD over 497 aligned residues. Both proteins originate from the same protein complex (PDB ID 8i1u). Ribbon representation (G), surface (H) and LBS (I) of a proteasome subunit β5 (orange, test ID 1625, test revealed ID 6epc_12:L:5_model1, class 20) and its Nearest Neighbor, a proteasome subunit β2 (green, train ID 8qyn_13_M_M_model1, class 69), discerned by the smallest Manhattan distance (λ_1_–λ_11_). This pair shares 7% sequence identity and 2.5 Å RMSD over 188 aligned residues. Purple vertical lines in C, F and I indicate differences between eigenvalues of the proteins being compared. The sum of these lines is the Manhattan distance (Equation 4).

The structural similarity was confirmed by conventional RMSD-based structure alignment, which resulted in 2.8 Å over 70 residues (Cα atoms) and a TM-score of 0.53 for histones (Figure 5A-C), where structural deviation was largely driven by a flexible loop in the N-terminal region; 1.9 Å over 497 residues and a TM-score of 0.89 for chaperonin subunits (Figure 5D-F); and 2.5 Å over 188 residues and a TM-score of 0.78 for proteasome subunits (Figure 5G-I). The TM-scores (range 0.0–1.0) of ≥0.5 indicated that the protein pairs shared the same fold (Xu and Zhang 2010).

To further contextualize these values and facilitate interpretation of the RMSD, which depends on both structural similarity and number of residues, we used fluorescent proteins GFP (PDB ID 1ema), YFP (PDB ID 7ugr; 90% sequence identity with 1ema; RMSD 0.4 Å; TM-score 0.99) and copGFP (PDB ID 2g3o; 19% sequence identify to 1ema; RMSD 1.8 Å; TM-score 0.85) (Figure S6) as references and the GFP:copGFP pair for normalization of the RMSD which was ∼0.01 Å per residue. The normalized RMSD values of the three pairs of proteins were 4.6, 0.4 and 1.5 for histones, chaperonin subunits and proteasome subunits respectively. This indicated that the supposedly misclassified (based on the SHREC protein classes) chaperonin subunits were structurally even more alike than the two fluorescent proteins based on conventional structure analysis whereas the two other pairs seemed still related.

Although TM-score, RMSD and the distance between the λ_1_–λ_11_ vectors use different approaches to capture protein structure similarity (atomic superimposition vs surface shape), their results were consistent, which, from our perspective, underlines the reliability of the LBS approach and, due to its rapid calculation, should facilitate the rapid structural classification of proteins. Curiously, the best structural alignment (2.5 Å RMSD) for the proteasome structures had only 7% sequence identity, whereas the sequence-based alignment (24% sequence identity) resulted in a much higher RMSD of 7.1 Å.

### 3.6 Analysis of suspected misclassifications II – functionally unrelated proteins

When assessing the 1-Nearest Neighbor distances of all test proteins assigned to non-homologous functionally unrelated classes (n = 21), we found that they were, on average, 4.5 times larger than correct classification distances, 2.8 times larger than homologous classification distances, and 2.1 times larger than non-homologous functionally related classification distances (Figure S9, Table S5). Three such non-homologous and functionally unrelated misclassifications were associated with large Manhattan distances to the 1-Nearest Neighbor of >0.005 in the whole training set, having therefore an even larger Manhattan distance to the closest training set protein in their own true class.

The largest of all 1-Nearest Neighbor distances in the whole SHREC 2025 dataset corresponds to a fragment of the 60S ribosomal protein L25 (test ID 1277, PDB ID 7r6q, chain H [auth X], class 62) and its 1-Nearest Neighbor found in class 78, corresponding to the SH3 domain of the c-Src tyrosine kinase (Figure S7). This misassignment may have occurred because test ID 1277 contained an additional N-terminal α-helix (Figure S7A) which was not present in most members of its class (162 of 169; bottom group of spectra in Figure S7L) and did not have any structural overlap with several members of its class (Figure S7E; 6 of 169). Alternatively, a spurious structural similarity between the C-terminus of the test structure (Figure S7A; test ID1277, PDB ID 7r6q) and that of the identified 1-Nearest Neighbor (Figure S7B; PDB ID 6c4s) might have resulted in some spectral similarity and hence the misclassification. Interestingly, eigenvalues λ_2_, λ_3_ and λ_4_ were almost perfect matches compared to a representative member of the class (PDB ID 6qt0, chain W [auth W]) indicating global structural similarity (Figure S7I). We think that this could be due to a coincidental shape similarity of the test shape N-terminus with the C-terminus of the training structure (which was missing the test shape; Figure S7 J and K). The second largest 1-Nearest Neighbor distance corresponded to another compromised mesh from a proteasome subunit B7 (test ID 1760, PDB ID 8qyj, chain K, class 71) with only 10% of residues modeled (Figure S4F). The third largest 1-Nearest Neighbor distance corresponded to a fully modeled ribosomal protein S21e (test ID 1513, PDB ID 4v6i, chain U [auth AT], class 83) which adopts a 3D structure different to all other members of its own class despite having an identical sequence. These examples illustrate the ability of the LBS to distinguish compromised surface meshes.

The rest of the misclassifications to non-homologous functionally unrelated classes (18 of 21) had Manhattan distances <0.005 to their 1-Nearest Neighbor in the training set, as observed for six test set surfaces from homologous classes 21, 64 and 94 of the SHREC 2025 dataset (Figure S8, Table S6). These classes represent members of the guanine nucleotide-binding protein Gα_i_1 family, which undergo major structural rearrangements during nucleotide exchange and interaction with regulatory partners, including the repositioning of helical domains (McClelland et al. 2020). This flexibility is reflected in experimental uncertainty about protein conformation and thus shape, resulting in a dispersion of the Laplace–Beltrami eigenvalues. Specifically, the average RSD of λ_1_–λ_11_ for surfaces in classes 21, 64 and 94 was 12.6% ± 7.4% compared to 2.1 ± 0.5% found for classes 54 and 57, corresponding to β-lactamase (Figure S10). Therefore, misclassifications for such proteins may arise if there are structurally similar (yet functionally unrelated) classes or densely populated regions in the descriptor space, where coincidental assignment of available surfaces into training and test sets can bring proteins belonging to different classes into accidental proximity. Accordingly, good coverage of possible surface conformations in the training set is necessary for the LBS approach to robustly classify protein surface shapes.

We found that such robust coverage was achieved for homologous classes 10 and 84, which contain influenza A virus polymerase basic protein 2 (PB2) surfaces. Like the guanine nucleotide-binding protein Gα_i_1 family above, the multidomain structure of PB2 can adopt different conformations based on experimental conditions and the sequence coverage of structures can vary across studies, for example, representing only the N-terminal domain (Keown et al. 2022; Krischuns et al. 2024). Importantly, the SHREC 2025 dataset contained 41 PB2 surfaces covering these different conformations and truncations, thereby ensuring they are well represented in the training dataset and hence facilitating a robust identification of test data based on the 1-Nearest Neighbor classification (Figure S11).

The likelihood of misclassification for the reasons discussed above can be minimized by adequate preparation of the training data, but there remains a small chance that visually dissimilar protein surfaces, in principle, exhibit small Manhattan distances between some of their LBO eigenvalues. This phenomenon is known as isospectrality (Reuter et al. 2006). Although it occurs rarely in 3D space, geometrically different surfaces become difficult to distinguish solely based on the LBS, especially if the protein surface exhibits conformational flexibility. For example, by visual inspection, we deemed some the 1-Nearest Neighbors identified for guanine nucleotide-binding protein Gα_i_1 family members to be distinct from the test structures to which they had been assigned (Figure S8; E vs G, I vs K, Q vs S).

### 3.7 Comparison with other shape retrieval methods

Fifteen protein classification methods were proposed for the SHREC 2025 dataset (Yacoub et al. 2025), grouped by the authors into three categories: 12 deep learning approaches, two conventional machine learning methods, and one training-free method (note that the difference between deep and conventional machine learning was not specified in the original publication). The latter implies that protein comparison and classification are performed without prior model fitting or parameter optimization on labeled training data. All but three (including the training-free approach) solely used the surface meshes as we did here. The classification performance was reported for the original 97 classes and a condensed list of 45 classes where homologous proteins had been grouped (Figure S10).

Among the surface-only methods, our LBS approach was surpassed only by a deep learning model comprising ∼5.7 million trainable parameters compared to the 11 values we use here (accuracy 91.9% vs 85.8%). LBS is therefore a computationally lightweight approach with feature extraction of eigenvalues λ_1_–λ_11_ requiring only 75 min on a standard CPU, whereas the deep learning approach required ∼470 min on dedicated GPU hardware. Moreover, model training required ∼23 h for the deep learning model but only ∼3 s for the 1-Nearest Neighbor assignments of our LBS method.

A training-free 1-Nearest Neighbor-based classification method similar to the LBS approach is 3D-SURFER. This is based on 3D Zernike polynomials and can be regarded as a compact representation of protein shapes by orthogonal series expansions on the unit sphere (Novotni and Klein 2003; Sael et al. 2008; Guzenko et al. 2020; Aderinwale et al. 2022). Normalization to a fixed spherical domain removes information about the surface scale, which can hinder the discrimination of proteins of different scales (e.g., small vs large globular proteins). Instead, electrostatic surface information split into positively charged, negatively charged and neutral areas is used, resulting in 363 descriptors. This introduces additional physicochemical complexity beyond pure geometry, which may complicate the interpretation of shape contributions and similarity, especially when seeking distant homologs. In contrast, LBO eigenvalues avoid the loss of native size information while capturing surface shape. Specifically, unlike 3D Zernike descriptors, we did not incorporate electrostatic features but deliberately focused on geometry alone in order to isolate the contribution of surface shape on classification performance.

Overall, 3D-SURFER was 2% more accurate when applied to the 45 homology-grouped classes (99.7% vs 97.7%), which corresponded to only eight misclassifications (Table S8), including the three test set proteins with the largest 1-Nearest Neighbor distances in our approach (Section 3.5). Likewise, one of the proteins we discussed in terms of experimental uncertainty (Figure 5) was also misclassified. Interestingly, 3D-SURFER failed to properly classify test ID 1882 (PDB ID 1ohh, chain G), corresponding to one ATP synthase γ subunit (class 38), which we classified correctly. It will be interesting to compare the performance of the two approaches when electrostatic properties are also incorporated into the LBS method. Using a single thread, our method reduced computational time by 96% compared with 3D-SURFER. Given the already low computational cost of our method, parallelization was deemed unnecessary, despite its use in the 3D-SURFER approach to reduce wall-clock time. Even when compared with their parallelized implementation, our method still achieved a 72% reduction in computational time. Nevertheless, if applied to databases with several thousand entries, our LBS approach is embarrassingly parallel which means the approach can readily be parallelized because each protein surface can be processed independently.

Beyond the classification approaches presented in the SHREC 2025 contest, several methods have been developed for large-scale protein structure comparison, including Foldseek and related frameworks designed for the rapid mining of expanding structural databases (van Kempen et al. 2024; Segura et al. 2026). Although highly efficient, these methods remain largely dependent on residue-level information, combining backbone-derived structural features with deep learning algorithms, which can limit the ability to identify structurally similar proteins with little sequence identity. In contrast, our LBS approach operates directly on protein surfaces and is entirely agnostic to sequence composition, residue identity, and prior training. By relying exclusively on the intrinsic geometry encoded in the LBS, it provides a purely surface-based and training-free alternative for structural comparison.

## 4. Conclusions

Using the first 11 eigenvalues of the LBS (λ_1_–λ_11_) and a training-free 1-Nearest Neighbor classification algorithm, we achieved a classification accuracy of 85.8% on the original 97 protein classes of the SHREC 2025 dataset and 97.7% when using the 45 homologous classes. We found that ∼1.4% of the misclassified proteins were similar in shape based on manual inspection and confirmation by atomic superimposition, leaving only 0.9% of proteins assigned to improper classes, which in some cases reflected compromised protein structures.

In the future, we will improve the performance of LBS by incorporating the targeted selection of eigenvalues from the spectrum based on the classification task, testing additional classification approaches beyond the 1-Nearest Neighbor method, and including confidence estimates from structure prediction methods, enabling uncertainty-aware shape representations. These modifications will help to disentangle genuine functional surface variability from experimental or modeling limitations, thereby improving robustness and interpretability in large-scale structural classification tasks. Detail improvements, such as using a dynamic probe size radius depending on protein surface hydrophobicity while generating the meshes, may increase accuracy (Halle and Davidovic 2003). Accordingly, our LBS approach will simplify protein structure comparison and may even form the basis of a new shape-driven rather than sequence-driven classification. The method may also be tailored to distinguish closely related groups of protein surfaces or to screen for distant homologs. Specifically, the sequence-independence of LBS classification can facilitate the identification of distant yet structurally and potentially functionally related proteins, for example to find enzyme variants for biotechnological applications. Additionally, other geometric descriptors derived from the LBS, like the heat kernel signature and wave kernel signature, as well as surface properties, such as electrostatic potential and hydrophobicity, can be used to improve classification even further. This may be directly relevant to biotechnological applications such as the prediction of protein–ligand interactions in chromatography or the identification of enzyme substrates.

## Declarations

### Ethics approval

Not applicable.

### Consent for publication

All authors have seen a draft version of the manuscript and concur with its submission to the journal.

### Authors’ contributions

BB and JF developed the concept. MF and JE designed the workflow. MF conducted the computational testing. MF and JB analyzed the data. MF prepared the figures and drafted the manuscript. JB revised the manuscript.

## Acknowledgements

We wish to thank Dr. Richard M Twyman for editorial assistance.

## Data availability

Data can be made available upon request to the corresponding author.

## Code availability

Code can be made available on request to the corresponding author.

## Abbreviations

CATH: Class, Architecture, Topology, Homologous superfamily
ECOD: Evolutionary Classification of Domains
MSMS: Michael Sanner Molecular Surface
PDB: Protein Data Bank
SCOP: Structural Classification of Domains

